# Correlation imputation in single cell RNA-seq using auxiliary information and ensemble learning

**DOI:** 10.1101/2020.09.03.282178

**Authors:** Luqin Gan, Giuseppe Vinci, Genevera I. Allen

**Affiliations:** Rice University; University of Notre Dame

**Keywords:** Single Cell RNA-seq, Imputation, Correlation Completion, Ensemble Learning, Auxiliary Information, Clustering, Dimension Reduction, Graphical modeling

## Abstract

Single cell RNA sequencing is a powerful technique that measures the gene expression of individual cells in a high throughput fashion. However, due to sequencing inefficiency, the data is unreliable due to dropout events, or technical artifacts where genes erroneously appear to have zero expression. Many data imputation methods have been proposed to alleviate this issue. Yet, effective imputation can be difficult and biased because the data is sparse and high-dimensional, resulting in major distortions in downstream analyses. In this paper, we propose a completely novel approach that imputes the gene-by-gene correlations rather than the data itself. We call this method SCENA: Single cell RNA-seq Correlation completion by ENsemble learning and Auxiliary information. The SCENA gene-by-gene correlation matrix estimate is obtained by model stacking of multiple imputed correlation matrices based on known auxiliary information about gene connections. In an extensive simulation study based on real scRNA-seq data, we demonstrate that SCENA not only accurately imputes gene correlations but also outperforms existing imputation approaches in downstream analyses such as dimension reduction, cell clustering, graphical model estimation.

## 1 Introduction

In genomics, researchers are interested in discovering the relationships between genes, monitoring changes of gene expression, and understanding the influence of genes on the organism. Bulk RNA sequencing (bulk RNA-seq) is a sequencing technology that lets us analyze gene expression from samples that contain a large number of cells by revealing the presence and quantity of RNA. With bulk RNA-seq data, significant results on gene-to-gene connection and gene-to-disease relationship can be obtained by machine learning methods, including dimension reduction, clustering models, and graphical models.

However, bulk RNA-seq only measures average gene expression levels across all cells in the sample, and it cannot detect gene expression differences across different types of cells. Single cell RNA sequencing (scRNA-seq) solves this problem by measuring the gene expression of individual cells, allowing us to clarify the critical difference among cells from the same organism. This genomic technology has helped discovering rare cells in different tissues by gene expression patterns and therefore is an important and powerful tool for transcriptome analysis [19].

Yet, data quality of scRNAseq is poorer than that of the bulk RNAseq, especially because of the presence of *dropouts*, technical artifacts where genes erroneously appear to have zero expression due to sequencing inefficiency. The loss of information is significant in scRNA-seq data, and can lead to major problems in downstream analyses.

Numerous *imputation* methods have been developed to fill in the dropout values in the scRNA-seq data. The SAVER model [14] predicts gene expressions under the assumption that the measured gene expressions follow Poisson-Gamma distributions, where the latent Gamma random variables are the true gene expressions. Based on the similarity among cells’ gene expressions, both drImpute [9] and PRIME [16] impute the dropouts of a cell by using the gene expressions of the cells belonging to the same cluster. The scRMD methodology [1] infers the gene expressions of cells by robust matrix decomposition, where the dropouts are encoded in a sparse matrix, and the matrix of true gene expressions is low rank. Other approaches include [30] and [37].

Yet, effective imputation of the missing values in scRNA-seq data can be difficult and biased because the data is sparse and high-dimensional – the number of genes is typically over 20,000 and the number of cells is usually only a few hundreds. In fact, researchers are interested in gene-to-gene connections and interactions, clustering of cells or principal component analysis of the scRNA-seq, and all these analyses require a well estimated correlation matrix of the gene expressions. Unfortunately, the presence of dropouts generates several challenges. For instance, if all zeros are assumed to be true values, the sample correlation matrix is corrupted. On the other hand, assuming that all zeros are dropouts, i.e. missing values, the Pearson correlation of two genes may be computed empirically only if there are enough pairwise-complete observations, otherwise it is infeasible. That is, the presence of dropouts can cause missingness in the sample covariance matrix. For this last problem there are plenty of covariance matrix completion methodologies that may be used [26, 21, 10, 17], but these, just like the data imputation methods [14, 9, 16, 1], perform ideally under different assumptions about the data.

However, in this challenging situation *auxiliary information* could be very helpful. Indeed, improvements in estimation performance due to the use of auxiliary information have been observed in various contexts [5, 11, 25, 8, 22, 31, 27, 23]. For instance, Hecker et al. [11] discuss improvements in network inference allowed by the incorporation of genome sequence and protein-DNA interaction data. Moreover, Lin et al. [25] study age-related macular degeneration by incorporating prior knowledge from previous linkage and association studies. Furthermore, Gao et al. [8] and Li et al. [22] use information about Gene Ontology annotation to improve network estimation. Finally, Novianti et al. [27] use gene pathway databases and genomic annotations to improve prediction accuracy, and Liang et al. [23] use auxiliary information about gene length and test statistics from microarray studies for the analysis of differential expression of genes.

In this paper we propose a novel approach, **SCENA** (*Single cell RNA-seq Correlation completion by ENsemble learning and Auxiliary information*), which estimates the gene-by-gene correlation matrix by incorporating auxiliary information about the underlying biological connections and other data sources together with the dropout-corrupted scRNA-seq data of interest. The auxiliary information we use includes the gene pathway database from the Kyoto Encyclopedia of Genes and Genomes (KEGG) [18], gene interaction networks from the Biological General Repository for Interaction Datasets (BioGRID) [28], protein-protein interaction networks from STRING [29], bulk cell RNA-seq data of 39 different tissues from the Encyclopedia of DNA Elements (ENCODE) [3], and other scRNA-seq data collected from cells of the same type of organism as the scRNA-seq dataset of interest. To implement SCENA, we first convert all these auxiliary data sources into a collection of correlation matrices, in addition to the correlation matrices recovered from scRNA-seq data via various imputation approaches and matrix completion strategies. Then, we ensemble all obtained correlation matrices into a final gene-by-gene correlation matrix estimate by *model stacking*.

We show that SCENA outperforms other existing methods in terms of correlation matrix completion, dimension reduction, clustering, and graphical modeling with an extensive simulation study (Section 3). Finally, we apply the methods to the analysis of a massive scRNA-seq data set of embryonic stem cells [2] (Section 4).

## 2 Method: SCENA

Let *X*_*s*_ be an *N* × *M* scRNA-seq data matrix of gene expressions of *N* genes measured over *M* cells. Because of technology limitations of the sequencing process, the data matrix *X*_*s*_ typically contains numerous false zeros called “dropouts”, where the transcript was not detected at the sequencing process. Thus, *X*_*s*_ must be seen as a corrupted version of some underlying data matrix *X* that is free from dropouts. To obtain estimates of the gene-by-gene correlations, in traditional approaches the data matrix *X*_*s*_ is typically first subject to some *imputation* process ℐ, which identifies the dropouts and predicts their values. The resulting imputed data matrix ℐ (*X*_*s*_) is then used to compute correlation estimates. Alternatively to this approach, we may complete or repair directly the correlation estimates obtained from *X*_*s*_ by standard *matrix completion* approaches, or approximate them by using various sources of *auxiliary information*.

Thus, several possible useful estimates of the single-cell gene-by-gene correlations are available, and a combination of them may let us obtain an ultimate reliable estimate of the gene-by-gene correlation matrix. Our proposed approach, **SCENA**, builds upon this strikingly simple but powerful idea of optimally combining multiple correlation matrix estimates. SCENA estimates the gene expression correlation matrix of scRNA-seq data by combining multiple genetic correlation matrices derived from various sources of information. In Section 2.1 we describe the derivation of several correlation matrix estimates, and in Section 2.2 we combine them via model stacking. In the rest of the paper, gene expressions are transformed according to *x* ↦ log_2_(1 + *x*) before computing correlations.

### 2.1 Single correlation matrix estimates

SCENA combines the following four groups of single correlation matrix estimates.

1. Blind correlation estimate (“all zeros are true”) This is the sample correlation matrix 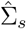 of the scRNA-seq data matrix *X*_*s*_ assuming all zeros are real zeros.
2. Imputation (“some zeros are dropouts and we try to correct them”). The imputation methods [14, 9, 16, 1] let us impute the scRNA-seq data, and thereby obtain correlation matrices.
3. Correlation matrices based on auxiliary data. Auxiliary is any kind of data that is beyond the scRNA-seq data of interest. We consider the following three kinds of auxiliary data, which can be used to compute correlation matrices.
  a. *Bulk RNA-seq data*. Given a matrix of bulk RNA-seq data, we calculate its sample correlation matrix. In this paper we use auxiliary bulk RNA-seq data of [3].
  b. *Other scRNA-seq data*. It is possible that another scRNA-seq data set 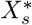 presents less dropouts for some of the genes in the main data matrix *X*_*s*_, so the lost information of those genes might be found in 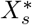. The sample correlation matrix of such additional data matrix 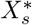 is computed. In this paper we use auxiliary scRNA-seq data from [35, 20].
  c. *Biological networks*. The KEGG pathway database [18] provides information about *n* = 6860 genes and *c* = 239 gene pathways which can be summarized in an *n* × *c* matrix *K* = [*K*_*ij*_] where *K*_*ij*_ = 1 if gene *i* is in pathway *j, K*_*ij*_ = 0 otherwise. We compute the *n* × *n* sample correlation matrix of *K*. From the BioGRID network [28] we extract an adjacency matrix *A* ∈ {0, 1}^*N* ×*N*^ of gene connections, and obtain the correlation matrix diag 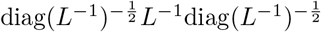, where *L* is the Laplacian matrix *L* = *D* − *A*, and *D* is a diagonal matrix with 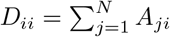, i.e. the degree of gene *i*. Finally, in the STRING network [29], we construct a correlation matrix by treating gene-by-gene combined connection scores as correlations.
4. Skeptical correlation estimates (“all zeros are dropouts”) Assuming all zeros are dropouts, we obtain the matrix 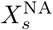 which corresponds to *X*_*s*_ with all zeros replaced by missing values NAs. From this matrix, it is possible to compute the *pairwise complete* sample correlation matrix 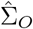, which contains missing entries for all those gene pairs with no jointly nonzero measured scRNA-seq expressions. We obtain completed versions of 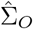 as follows.
  a. *Matrix completion*. We use [26] to produce a complete correlation matrix.
  b. *Convex combinations with matrices in (3)*. Given a correlation matrix derived from auxiliary data, say 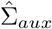, we can obtain a completed version of the pairwise complete scRNA-seq sample correlation matrix 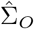 as 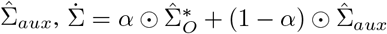, where 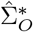 is a version of 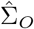 with all NA’s replaced by zeros, and *α* = [*α*_*ij*_] is a *N* × *N* weight matrix. We consider two types of weights:
    i. *Simple replacement*

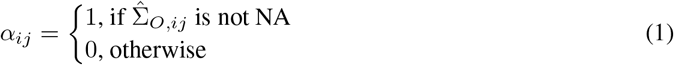
    ii. *Signal-to-noise ratio*

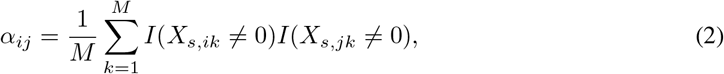

which is the proportion of cells where genes *i* and *j* have jointly nonzero read counts.

### 2.2 Model stacking

Let 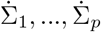 be the single correlation matrix estimates derived in Section 2.1. We aim to obtain a final correlation matrix estimate 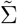 by model stacking in the form

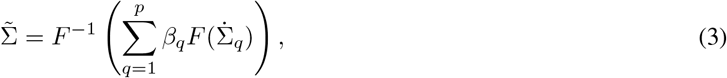

where *F*: ℝ^*N* ×*N*^ → ℝ^*N* ×*N*^ is an invertible mapping, and *β*_1_, …, *β*_*q*_ ∈ ℝ. The simplest choice of *F* is the identity mapping *F* (*A*) = *A*, ∀*A* ∈ ℝ^*N* ×*N*^, which however does not guarantee 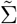 to be a positive semi-definite correlation matrix, even if all 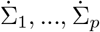 are positive semi-definite correlation matrices, unless we impose appropriate constraints on *β*_1_, …, *β*_*q*_. For instance, a sufficient condition is *β*_*q*_ *>* 0, ∀*q*, with Σ_*q*_ *β*_*q*_ = 1, which specifies a convex linear combination. Another possible mapping is *F* (*A*) = *A*^−1^, which requires 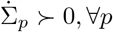. In any case, if 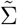 is not a positive semi-definite correlation matrix, we replace it with the nearest correlation matrix [13] as per 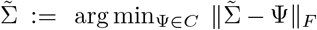, where *C* is the set of positive semi-definite correlation matrices.

There are many possible ways to specify Equation (3). We consider the following ones.

#### 2.2.1 Simple average

The simple average is obtained by setting *F* = identity mapping and 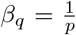, for all *q*, yielding the convex linear combination

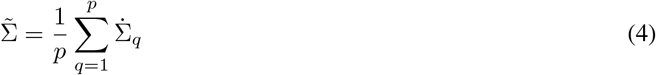

Since the weights are prespecified, this approach requires no additional tuning or validation steps. Also, if 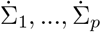 are all positive semi-definite correlation matrices, so is 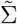. We will denote this solution by **SCENA**_**average**_.

#### 2.2.2 Regression

We assume a linear relationship between the true underlying correlation and the single correlation estimates,

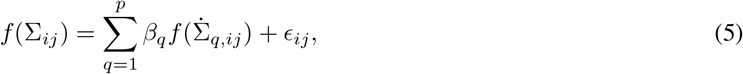

where *f*: (−1, 1) → ℝ is an invertible function, e.g. the Fisher transformation 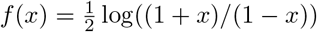, and *ϵ*_*ij*_ is an error component. The vector of coefficients *β* = (*β*_1_, …, *β*_*q*_)^*T*^ is then estimated by solving the penalized optimization problem

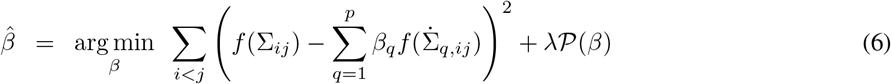

where 𝒫 is a penalty and *λ* ≥ 0 is selected via cross-validation. Cross-validation lets us reduce the risk of overfitting, and is implemented by creating multiple held-out data subsets that are iteratively removed from training and used instead to validate prediction accuracy. Setting 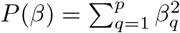 produces the *ridge estimator*, and the resulting final correlation matrix 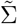 which will be denoted by **SCENA**_**ridge**_.

Of course, we do not know Σ_*ij*_ in Equation (5), but we can identify a small subset of genes and cells which we may assume to contain very few dropouts and could give us reliable estimates of Σ_*ij*_. Thus, to fit the regression model in Equation (5), we first extract a reference data matrix *Y*′ from the scRNA-seq data matrix *X*_*s*_ (Algorithm A), then compute the sample correlation matrix 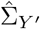 of *Y*′, and finally extract the off-diagonal entries which will be used as the response vector of the regression. Then, we obtain multiple perturbed versions of *Y*′ by creating artificial dropouts (Algorithm A). The off-diagonal entries of the matrices 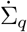, for *q* = 1…*p*, based on the perturbed data, are used as predictors. Algorithm A summarizes the full procedure.

## 3 Simulations

In this section we present an extensive simulation study showing that SCENA is superior to other methods in terms of correlation matrix completion, dimensionality reduction, clustering, and conditional dependence graphical modeling. In Section 3.1 we describe how we generate realistic artificial scRNA-seq data based on real data sets. Specifically, given a real scRNA-seq data set, we first extract a *reference data set*, a subset of data where all zeros can be safely assumed to be true values and not dropouts. Then, we generate *downsampled data* by creating dropouts in the reference scRNA-seq data according to the Poisson-Gamma scheme in Algorithm A. Finally, in Section 3.2 we assess the performance of SCENA and other existing methods at recovering the correlation structure of the reference data based on the corrupted downsampled data and other available auxiliary data.

### 3.1 Generating scRNA-seq data

#### 3.1.1 Original data sets

We use three human scRNA-seq data sets in this simulation study:

1. **chu**: human embryonic stem cells [2].
2. **chu_time**: human definitive endoderm cells (time-series sequencing) [2].
3. **daramanis**: human brain cells [4].

The number of cells and number of cell types are reported in Table 1 (genes with zero expression in all cells are removed).

**Table 1:** Human scRNA-seq data sets used in simulations.

|  | **chu** | **chu_time** | **darmanis** |
| --- | --- | --- | --- |
| **Tissue** | embryonic stem cells | definitive endoderm cell | brain cell |
| **# cell types** | 7 | 6 | 5 |
| **# cells** | 1,018 | 758 | 366 |
| **# genes** | 21,413 | 18,294 | 17,738 |
| **% zeros** | 47.43% | 51.15% | 80.06% |
| **# reference cells** | 951 | 689 | 332 |
| **# reference genes** | 2,522 | 2,160 | 2,306 |
| **citation** | [2] | [2] | [4] |
| **GEO accession code** | GSE75748 | GSE75748 | GSE67835 |

#### 3.1.2 Reference data sets

For each of the three data sets, we first match the genes with those available in the auxiliary data (Section 2.1), and then apply Algorithm A to perform quality control by filtering out low quality genes and cells, and finally extract reference data. All values in the reference data are treated as true gene expressions, i.e. the reference data is free from dropouts. The dimensions of the three resulting reference data sets are reported in Table 1.

#### 3.1.3 Downsampled data

For each of the three reference data sets, we apply Algorithm A to generate downsampled versions of the reference data. We set *s* = 10, and *r* = 3000, 1000, 1000 for chu, chu_time and darmanis data, respectively, to ensure the expected percentage of zeros in the downsampled data to be similar to the percentage of zeros in the original scRNA-seq data (Table 1).

### 3.2 Models comparison

In this section we assess the performance of SCENA and other existing methods at recovering the correlation structure of the reference data based on the corrupted downsampled data and other available auxiliary data. We show that SCENA_average_ and SCENA_ridge_ (Section 2.2) outperform SAVER, drImpute, scRMD, and PRIME in terms of correlation matrix completion, dimension reduction, clustering, and graphical modeling. The results shown are averaged across multiple downsampled data sets.

#### 3.2.1 Correlation completion

We measure the similarity between the correlation matrix estimators considered and the reference correlation matrix 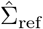 in terms of *mean squared error* (MSE) and average *correlation matrix distance* (CMD) [12]. Table 2 shows that SCENA outperforms all other data imputation methods in terms of both MSE and CMD. The baseline is set to be the MSE and CMD between 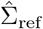 and the sample correlation of the downsampled data (blind estimate 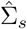, Section 2.1), treating all 0s as true gene expressions. SCENA_ridge_ has the lowest MSE and CMD among all methods as well as the baseline in chu_time and darmanis data. SCENA_average_ is also better in correlation completion than SAVER and PRIME.

**Table 2:**
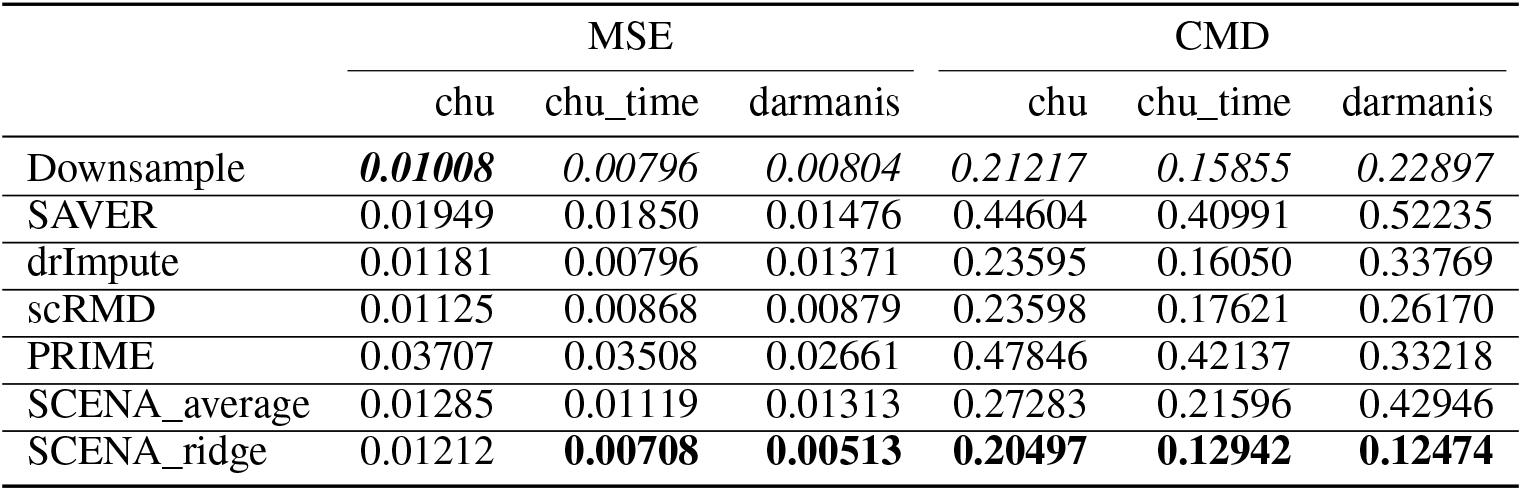
Correlation completion accuracy. MSE and Correlation Matrix Distance (CMD) between the reference correlation and the estimated correlation derived from various methods. SCENA_ridge_ has the lowest MSE and CMD among imputation methods in chu_time and darmanis data, and is lower than the baseline (“Downsample”), which is the sample correlation matrix of the downsampled data. All other imputation approaches perform worse than the baseline.

#### 3.2.2 Dimension reduction

For each correlation matrix estimate from inputed data and SCENA, we compute the matrix of eigenvectors *V*, and obtain the principal component scores *U* = *Z*^*T*^ *V*, where *Z* is a standardized version of the log transformed scRNA-seq data *X*_*s*_. In Figure 1, we compare the scatterplots of the top two PC scores of the cells against each other. The plots are colored by cell type labels, and PCs are derived from sample correlations of reference data, SAVER imputed data, and correlation estimations of SCENA_ridge_ and SCENA_average_. Both SCENA_ridge_ and SCENA_average_ recover the reference data structure better than data imputation, and yield scatterplots with a clear separation among different types of cells indicated by the cell type labels.

**Figure 1:**
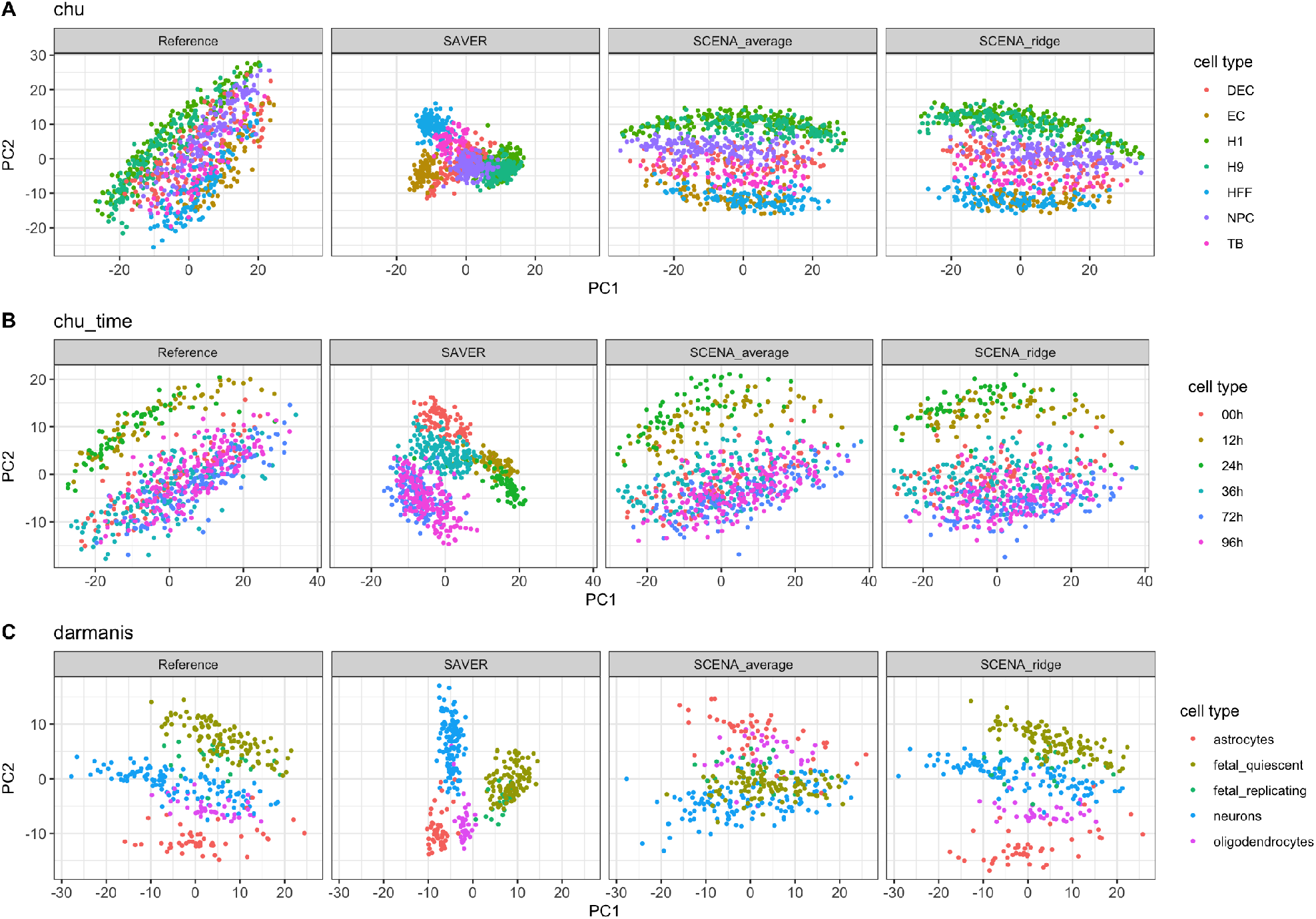
Dimension reduction accuracy. Scatterplots of the top two PC scores of the cells colored by cell type. Both SCENA_ridge_ and SCENA_average_ appear to recover the reference data structure better than SAVER, yielding scatterplots with a clear separation among different types of cells.

#### 3.2.3 Clustering

We perform hierarchical cells clustering (Ward’s minimum variance method with Manhattan distance; [7]) based on the standardized principal components of the downsample scRNA-seq data obtained from the different approaches considered (quantities *U* computed in Section 3.2.2). For each method, we use the top PCs with proportion of variance explained within the range (90%,99%), and set the number of clusters equal to the number of true cell labels in the scRNA-seq data. To assess clustering performance, we measure the similarity between cluster assignments and true cell labels by calculating the *adjusted rand index* (ARI). This metric takes values in the interval [0, 1], with large values indicating stronger similarity. In Figure 2 we can see that SCENA_average_ yields the best clustering performance over all other methods in all three datasets. Interestingly, in the chu data, SCENA_average_ has even better performance than the clustering obtained from reference data, in accordance with the fact that SCENA exploits auxiliary information besides the scRNA-seq data.

**Figure 2:**
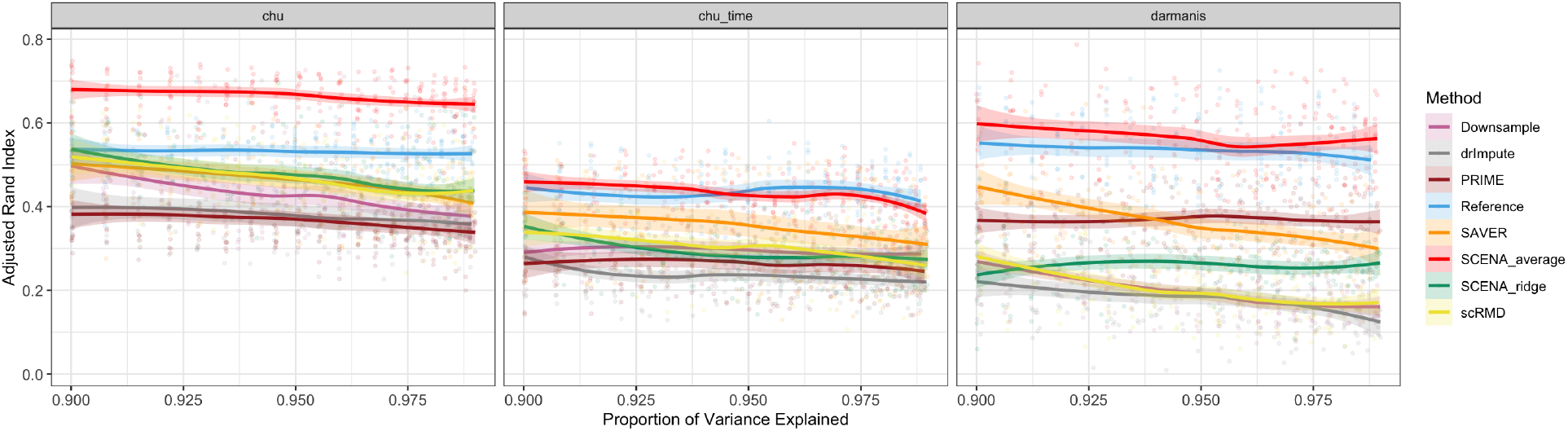
Clustering performance. Adjusted rand index (higher is better) of cell type grouping via hierarchical clustering after dimension reduction via PCA explaining various proportions of variance. SCENA_average_ yields the best clustering performance over all other methods in all data sets, and even better than the clustering obtained from the reference data in the chu data.

#### 3.2.4 Conditional dependence graphs

Conditional dependence graph estimation in the case where several pairs of variables are never observed jointly is a major statistical problem that has gained strong interest recently; a thorough theoretical investigation of the so called *graph quilting problem* can be found in [32]. Such problem is strictly related to ours, where an extremely large number of gene pairs have no reliable empirical correlation estimates. Here, we investigate the graph recovery performance based on the various correlation matrix estimates via simulations. Specifically, we plug the correlation matrix estimates into the graphical lasso [36] to obtain gene-by-gene conditional dependence graphs via sparse precision matrix estimation. For simplicity, we compute graphs about only the top 50 most variable genes among cell types, identified by applying ANOVA to gene expression of reference data adjusted by cells’ library sizes. To evaluate the graph recovery performance of a method, we compute the F1-score with respect to the graph estimated from reference data. In Figure 3A we plot F1-score versus number of graph edges for all methods and data sets. SCENA_ridge_ is superior in recovering the reference graph than other methods in chu and darmanis data, and it produces a similarly high F1-score in chu_time data as SAVER method. For illustration, in Figure 3B we also display conditional dependence graphs relative to the chu data with 50 edges.

**Figure 3:**
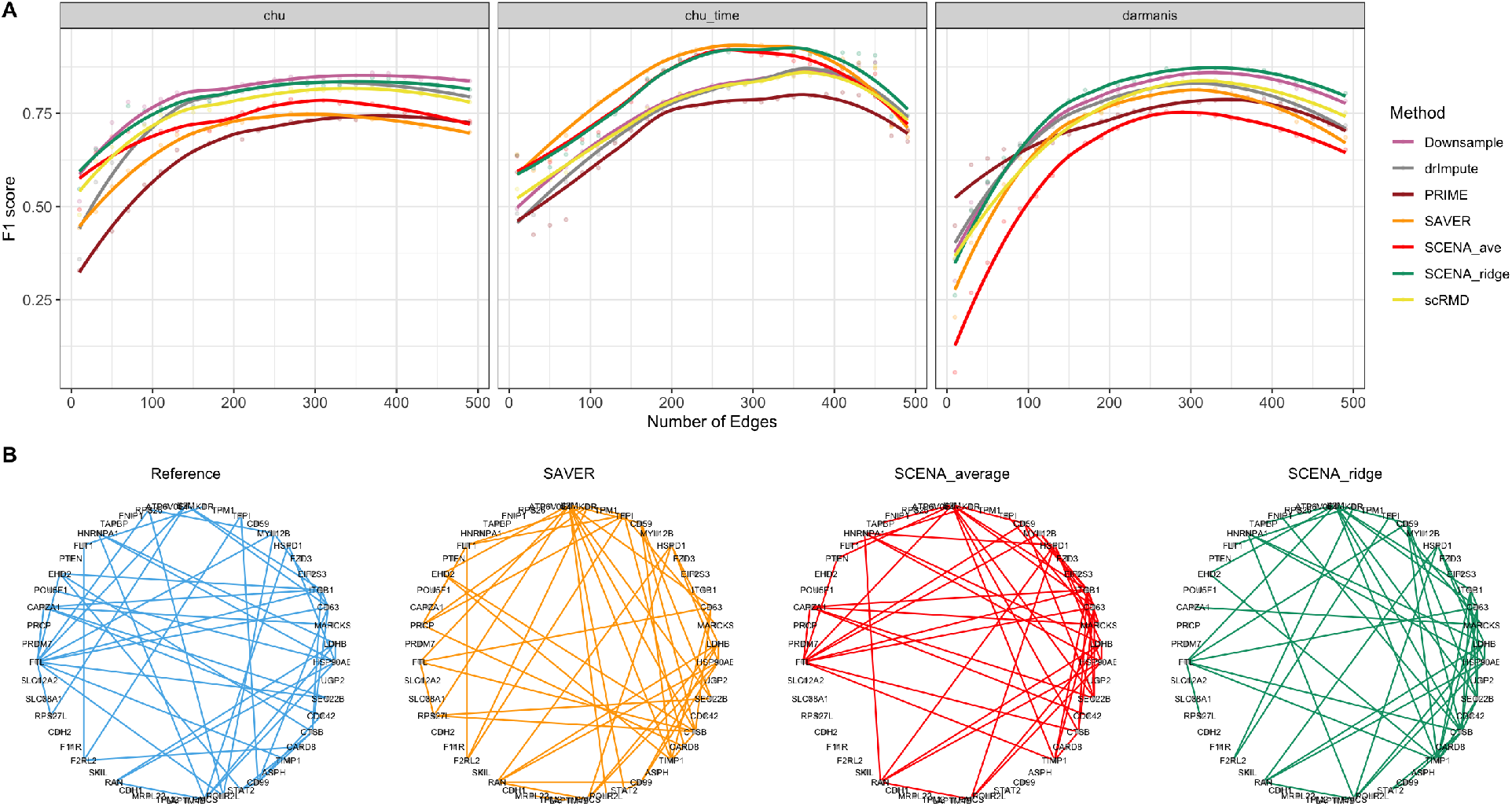
Genetic graph recovery. A: F1 score (higher is better) quantifying the performance of methods at recovering the reference conditional dependence graphs of 50 most variable genes for various numbers of edges. SCENA_ridge_ exhibits strong performance for all data sets, while other methods’ performance dramatically changes across different data sets. B: Conditional dependence graphs of chu data, setting the number of edges to 50.

## 4 Application to stem cell data

We now apply the methods to the analysis of the chu data set (Table 1) containing the gene expression of 6,038 genes (largest genes set that matched available auxiliary information) measured in 1,018 human embryonic stem cells. In Figure 4A we plot the first two principal components based on SCENA_average_ and SCENA_ridge_, while in Figure 4B we compare the cell clustering performance of SCENA with other methods in terms of ARI. The hierarchical clustering based on SCENA_average_ performs the best at recovering true cell type labels, in accordance with simulation results (Section 3.2.3). Finally, in Figure 4C we display the conditional dependence graph (graphical lasso; [36]) of the 30 most variable genes among cell types (ANOVA criterion as in Section 3.2.4) based on SCENA_ridge_, with number of edges 163 selected via Extended Bayesian Information Criterion (EBIC, [6]). The protein coding gene *DNMT3B* is the hub node with largest number of connections (20 edges). This result is reasonable because *DNMT3B* is a catalytically active DNA methyltransferase [24], and is specifically expressed in totipotent embryonic stem cells [34]. Moreover, *DNMT3B* is one of the pluripotency markers with high level of expression in the cell type H1, as demonstrated by [2]. Besides, genes *IFI16* [15] and *HAND1* [33] are marker genes in cell type EC and cell type TB, respectively, and correspondingly have relatively large numbers of connections.

**Figure 4:**
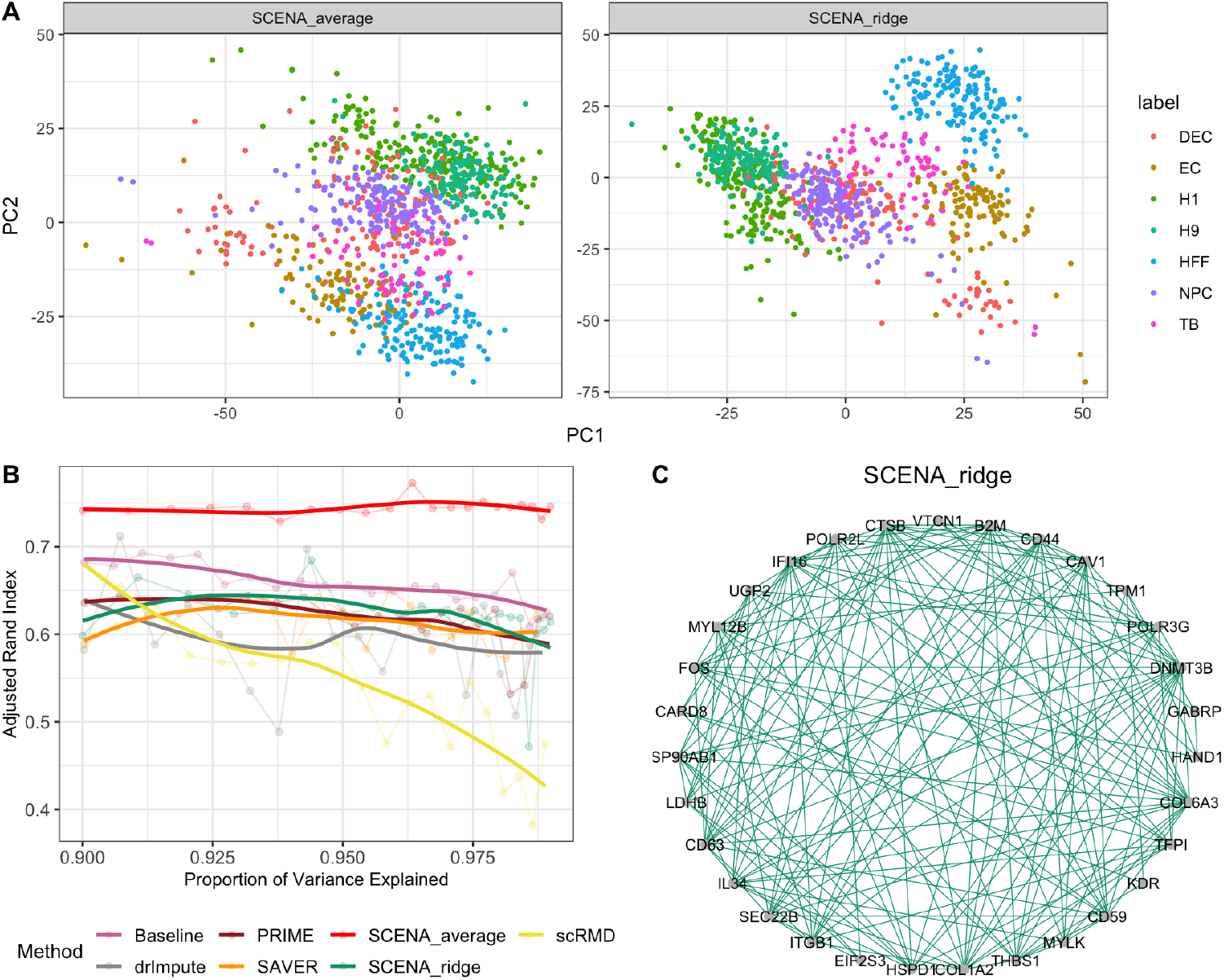
Real data application results for the chu data. A. Dimension reduction: scatterplots of the top two PC scores of the cells colored by cell type. B. Clustering: adjusted rand index of cell type grouping via hierarchical clustering after dimension reduction using PCA explaining various proportions of variance. SCENA_average_ yields the best clustering performance over all other methods. C: Conditional dependence graph (graphical lasso; 163 edges selected via EBIC) based on SCENA_ridge_ correlation estimate. Gene *DNMT3B* is the hub node with the largest number of connections (20 edges). This result is supported by the scientific literature as *DNMT3B* is one of the pluripotency markers with high level of expression in the cell type H1 [2]. Also, genes *IFI16* and *HAND1* are marker genes in cell type EC and cell type TB, respectively, and correspondingly have relatively large numbers of connections.

## 5 Discussion

We have proposed and studied SCENA, a novel methodology for gene-by-gene correlation matrix estimation from dropout-corrupted single cell RNA-seq data. SCENA builds upon the strikingly simple but powerful idea of optimally combining multiple gene-by-gene correlation matrices derived from various sources of information, besides the scRNA-seq data of interest. This combination is implemented efficiently via model stacking techniques.

We have demonstrated that SCENA can provide superior estimation performance compared to traditional data imputation methods. In our analyses, SCENA_ridge_ remarkably recovered the information underlying the corrupted scRNA-seq data in terms of correlation completion, dimension reduction, and graphical modeling, while the hierarchical clustering based on SCENA_average_ yielded cell groupings that best reflected true cell type heterogeneity in terms of adjusted rand index. Indeed, although both variants combine the same single correlation matrices via model stacking, the weighting coefficients of SCENA_ridge_ are calibrated for the optimal recovery of the true correlation structure of the corrupted gene expression data, while SCENA_average_ simply assigns uniform weights, presumably upweighting auxiliary biological network structures that are more informative about cell characteristics. SCENA_average_ is computationally cheaper than SCENA_ridge_, because the estimation of the weighting coefficients of SCENA_ridge_ involves multiple additional imputation and optimization steps that are computationally expensive. For instance, in the application presented in Section 4, the model stacking step for SCENA_ridge_ took about 40 minutes, while only about 30 seconds for SCENA_average_, on a laptop with 16GB of RAM (2133 MHz) and dual-core processor (3.1 GHz). Given all these considerations, we recommend to use SCENA_average_ for the analysis of massive scRNA-seq data sets.

While we have demonstrated our approach using specific auxiliary sources, SCENA is general and conducive to many different types of correlation imputation approaches and additional sources of auxiliary information on genetic interactions. Additionally, our approach can be further optimized using different machine learning approaches to model stacking and ensemble learning. Overall, we expect SCENA to become an important instrument for downstream analyses of massive scRNA-seq data that powerfully incorporates known auxiliary information on genetic interactions.

L.G., G.V., and G.A. are supported by NIH 1R01GM140468, NSF DMS-1554821 and NSF NeuroNex-1707400. G.V. is additionally supported by a Rice Academy Postdoctoral Fellowship and the Dan L. Duncan Foundation.

## A Algorithms

[Reference data selection]

Input: *N* × *M* data matrix *X*; parameter vector *a*.

1. Filter out cells with library size greater than *a*_1_-th percentile.
2. Remove genes with mean expression less than *a*_2_-th percentile.
3. Remove genes with less than *a*_3_-th percentile non-zero cells.
4. Keep cells with library size greater than the *a*_4_-th percentile.
5. Keep genes with non-zero proportion greater than *a*_5_-th percentile.

Output: *N*′ × *M*′ reference data matrix *Y*. We use default values *a*_1_ = 95, *a*_2_ = 25, *a*_3_ = 15, *a*_4_ = 5, *a*_5_ = 50.

[Poisson-Gamma downsampling]

Input: *N* × *M* data matrix *X*; parameters *s, r >* 0.

1. Draw 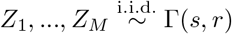.
2. Draw 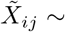 Poisson(*X*_*ij*_*Z*_*j*_), for *i* = 1, …, *N* and *j* = 1, …, *M*.

Output: *N* × *M* downsampled data matrix 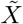.

[Stacking regression validation]

Input: *N* × *M* scRNA-seq data matrix *X*; reference parameter vector *a*; downsampling parameters *s, r >* 0; collection of *N* × *N* single correlation matrices (Section 2.1); number of downsampling repeats *B*; transform function *f*.

1. Obtain reference *N*′ × *M*′ data matrix *X*′ via Algorithm A with parameter *a*.
2. Construct response vector 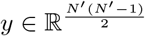 by extracting off-diagonal entries from 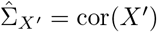.
3. For *b* = 1, …, *B*:
  a. Generate downsampled *N*′ × *M*′ data matrix 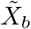 from *X* via Algorithm A with parameters *s, r*.
  b. Obtain all *N*′ × *N*′ single correlation estimates based on 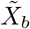 and auxiliary correlation matrices 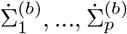.
  c. Construct predictors matrix 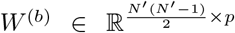 by extracting off-diagonal entries from each 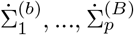.
4. Compute 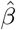 by regressing *f* (**y**) on *f* (**W**) via Equation (6), where **y** = (*y*^*T*^, *y*^*T*^, …, *y*^*T*^)^*T*^ and **W** = (*W* ^(1)*T*^, …, *W* ^(*B*)*T*^)^*T*^.

Output: *N* × *M* correlation matrix 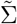 via Equation (3) with vector of coefficients 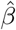.

